# Neural Mechanisms of Mixed Speech and Grasp Representation in Sensorimotor Cortices

**DOI:** 10.64898/2026.01.02.697428

**Authors:** Crispin Foli, Emily C. Conlan, William D. Memberg, Preethisiri Bhat, Emily L. Graczyk, Tyler R. Johnson, Dawn M. Taylor, Eric Z. Herring, Jennifer A. Sweet, A. Bolu Ajiboye

## Abstract

Recent brain-machine interface (BMI) studies have challenged traditional views of functional specialization in human motor cortices, suggesting that regions associated with hand control also support speech. The extent of this dual functionality as well as the neural mechanisms underlying it are unclear. We address this by analyzing intracortical neural activity from seven brain regions (spanning motor, premotor, somatosensory and parietal regions) across two human participants with tetraplegia. Across all regions, grasp decoding was robust. In addition, we achieved reliable discrete-word decoding during silent reading as well as vocalized speech. Both tasks largely recruited overlapping neural populations within each region, yet these populations reconfigured their functional connectivity between tasks. Additionally, subspace analyses revealed segregated computations for speech and grasping despite mixed selectivity at the single channel level. Our findings support multi-functional BMIs capable of decoding speech and grasping from the same implant and highlight ventral premotor area 6r as a novel target.

## Introduction

Advances in intracortical brain-machine interfaces (BMIs) have expanded our understanding of the neural basis of motor control, offering significant potential for improving the efficacy of BMI-based rehabilitative interventions for individuals with chronic paralysis (1–4). These BMI technologies have traditionally targeted the “hand knob” area of precentral gyrus because of the conventional belief in the somatotopic and segmented organization of the motor strip (5). However, the brain’s functional architecture, particularly within motor areas such as the precentral gyrus may not be as localized as previously assumed. Historically, subregions such as the ventral premotor cortex (PMv) and primary motor cortex (M1), have been viewed primarily through the lens of their distinct roles in motor planning and execution. Several human studies, based on functional imaging and transcranial magnetic stimulation (TMS), have consistently implicated PMv in motor planning related to hand–object interactions (6–9). Specifically, PMv has been identified as a critical region for shaping hand posture during grasping. Causal evidence from TMS studies shows that transient disruption of PMv impairs precision grip execution (7), particularly the fine finger control needed for accurate object contact (6). Moreover, PMv exhibits grasp-specific interactions with M1, exerting a net-inhibitory influence at rest that is released or converted to facilitation during motor execution depending on grasp type (7). PMv also contributes to predictive force scaling during grasping (6), further underscoring its central role in motor coordination.

However, emerging evidence from recent intracortical BMI studies have challenged the prevailing notion of functional specialization in putative motor areas, revealing that brain regions associated with hand motor function also exhibit selective tuning to speech (10–15). For example, single units in nominal “hand knob” area of M1 have been found to encode spoken syllables (10, 11), non-speaking orofacial movements (11) and imagined handwriting of individual letters and symbols (12). Multi-unit neural populations in PMv (area 6v) across multiple participants have been shown to encode both inner speech (i.e., a participant’s internal monologue) and attempted speech as well as different categories of non-speaking orofacial movements (13, 15, 16). Additionally, in a dual-task study, Wandelt et al. (14) successfully decoded imagined grasps from primary somatosensory cortex (S1). They also found that single units in the supramarginal gyrus (SMG), a posterior parietal cortex subregion previously linked to tool use, encode both silent and overt speech in addition to grasp type.

Collectively, these studies demonstrate the deviation of multiple brain regions from their traditionally ascribed functions and highlight the co-existence of motor and language representations within certain brain regions, blurring the boundaries between these two functions.

It is unclear if mixed selectivity for speech and grasping is a generalizable property of sensorimotor cortical regions. Additionally, the neural mechanisms allowing the dual representation of speech and grasping in cortical areas classically implicated in motor control are unclear. For instance, is multi-task representation within individual brain areas subserved by non-overlapping, task specialized neural clusters, or do different tasks recruit the same neural populations? We propose that the dual representation of language and motor functions within individual cortical regions is facilitated by *mixed selective* neural populations that reconfigure their functional architecture to perform different computations between tasks. We refer to individual neural elements (i.e., single channels etc.) as *mixed selective* if they are capable of encoding more than one task type or variable (17).

In this study, we present findings from ten intracortical implants (across two human participants) spanning the primary motor cortex (M1), primary somatosensory cortex (S1), parietal cortex (anterior intraparietal area, AIP), frontal cortex (inferior frontal gyrus, IFG), and ventral premotor cortex (PMv) (Fig. 1a). The two participants, RP1 and RP2, were enrolled as part of the Reconnecting the Hand and Arm to the Brain (ReHAB) clinical trial, which aims to restore sensation and hand motor control to persons with chronic cervical spinal cord injury (18). For this work, we place particular emphasis on two subregions of PMv: area 6v and its anterior–inferior counterpart, area 6r. The role of area 6v (often referred to interchangeably with “PMv” in the literature) is well established in both speech processing (13, 15, 16) and hand motor control (6–9, 14, 16). In contrast, the function of 6r remains less characterized. Anatomically, 6r lies between two functionally distinct regions, sharing its anterior border with the speech-dominant area 44 (part of Broca’s area) and its posterior–superior border with 6v (19). As a result, it has been dubbed a “transitional” region, bridging nominal motor and language cortices (20). Given its microstructural similarity to 6v (20) and its established structural (19) and functional (21) connectivity with classical motor areas, we hypothesized that 6r, like 6v, would support selective activation during grasping as well as speech.

**Figure 1.**
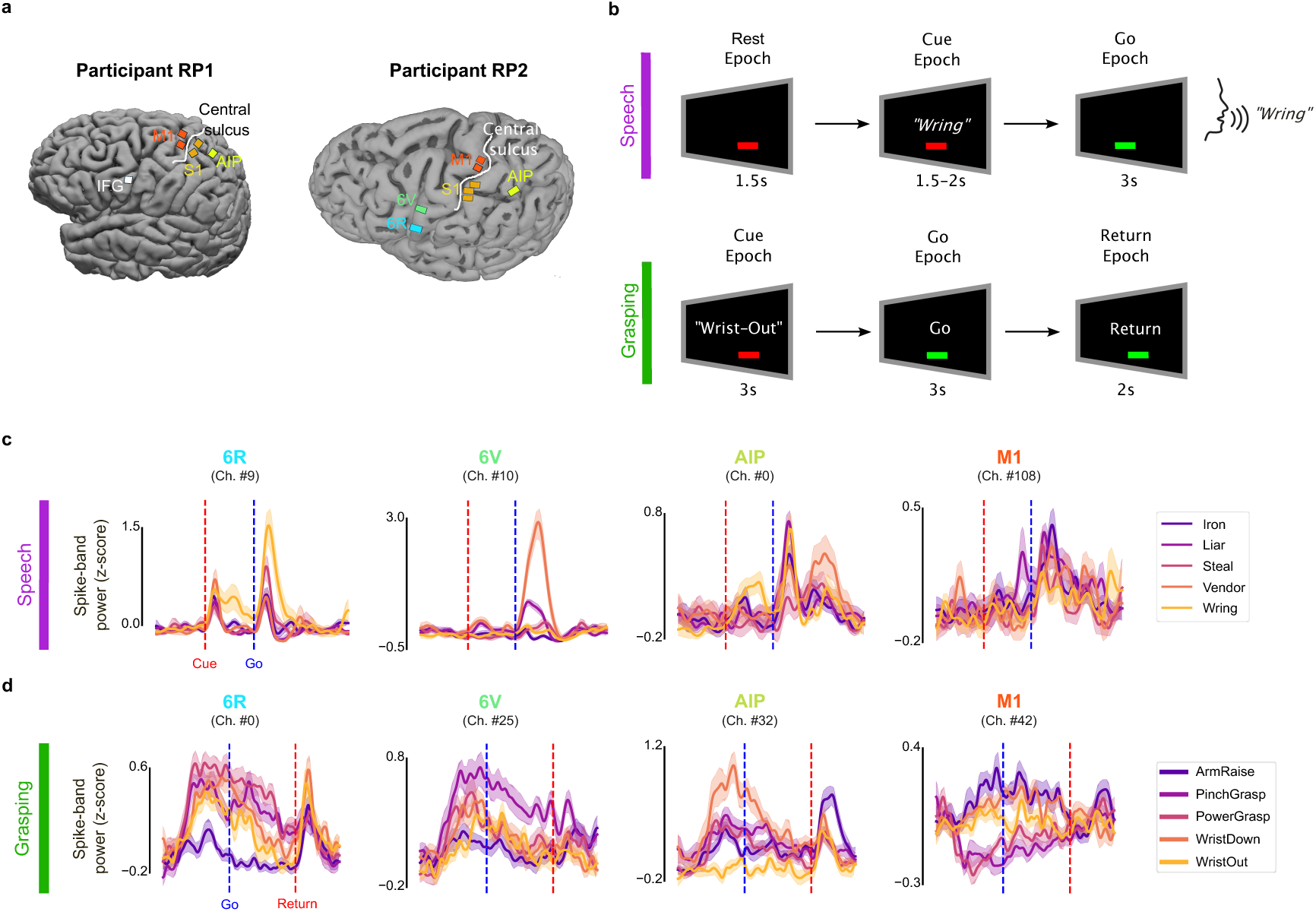
Experimental outline and neural representation of speech and grasping. **(a).** Illustrations of microelectrode array implant locations. Participants RP1 and RP2 were implanted as part of the ReHAB clinical trial (see methods), targeting AIP, M1, and S1 in both participants as well as IFG in RP1, and 6r and 6v in RP2**. (b).** Experimental plan for the speech and grasp tasks. The speech task (top) involved silent reading, followed by overt articulation of a cued word. A total of five unique words were cued. The motor task (bottom) involved attempted execution of a cued hand movement. This study focuses on five right hand movements. (RP1 performed slightly modified versions of these tasks which are illustrated in supplementary Fig 1). **(c, d).** Example neural features showing robust tuning to speech **(c)** and grasping **(d)** across 6r, 6v, AIP and M1 in RP2. The features displayed are z-scored spike-band power activity smoothed with a Gaussian kernel (standard deviation: 80ms, for visualization only). *(AIP–anterior intraparietal area; 6r–ventral premotor cortex area 6r; 6v–ventral premotor cortex area 6v; S1– hand area of primary sensory cortex; M1–hand area of primary motor cortex; IFG–Inferior Frontal Gyrus)*

AIP is traditionally implicated in motor planning, and contains visual and visuomotor neurons tuned to object features such as shape, size, and graspable surfaces (22–24). IFG, contains visuomotor and motor neurons that encode grasp type (22, 25, 26).

Multiunit neural activity across the cortical regions specified above were analyzed during speech and motor experiments with three primary objectives. First, we sought to determine whether mixed representations of both speech and grasping within the same cortical areas (which have previously been seen predominantly in area 6v (16)) generalize across multiple, functionally distinct cortical networks. Second, we sought to elucidate the role of area 6r (a region whose function in the human brain remains poorly understood) by examining its contributions to both motor and language encoding. Third, we aimed to identify key neural mechanisms of information representation that enable mixed speech and grasp representation, focusing in particular on neural subspace segregation. By addressing these aims, this study advances understanding of dual-task representation in the human cortex and informs the design of multifunctional BMIs capable of integrating communication and motor control.

## Results

### Both speech and grasping are represented within each brain area

Broadband signals were recorded while participants sequentially performed speech and grasping tasks (Fig. 1b, supplementary Fig. 1). In the grasping task, participants attempted up to five hand movements using an instructed-delay paradigm (Fig. 1b, bottom). The speech task involved silently reading a randomly cued word and then vocalizing it during the go epoch (Fig. 1b, top). Participant RP1 performed a slightly modified version of this task involving grammatical inflection (i.e., verb conjugation) of up to 47 different words (supplementary Fig. 1). Neural features across example electrodes showing robust, task-specific modulation of neural activity are shown in Fig. **1c–d** for different brain regions. PMv implants revealed a variety of rich electrode-level tuning to both vocalized and silent speech (i.e., during silent reading epoch). Notably, we identified individual electrodes showing preferential activation during vocalized speech only as well as those showing strong activation across both vocalized and silent speech (supplementary Fig. 2).

All subsequent RP2 analyses used windowed features (a 2s window preceding the go cue for grasping and a 1.5s window during the go epoch for speech). In order to quantitatively assess how well the neural populations in each area represented each task, we trained non-linear support vector machine (SVM) classifiers to predict task conditions from windowed features across an optimally selected electrode subset (see Methods, supplementary Fig. 3). We obtained performant results across multiple areas during silent reading, vocalized speech and grasp decoding (Fig. 2a) in participant RP2. Results in RP1 involved binary kinematic (predicting whether hand was opening or closing) and speech decoding (distinguishing covert verb conjugation versus overt articulation from neural activity). Ten-fold cross-validation was applied across all decoding analyses to mitigate bias. The resulting confusion matrices from the predictive analyses described above are shown in Fig. **2** **(b, c)**.

**Figure 2.**
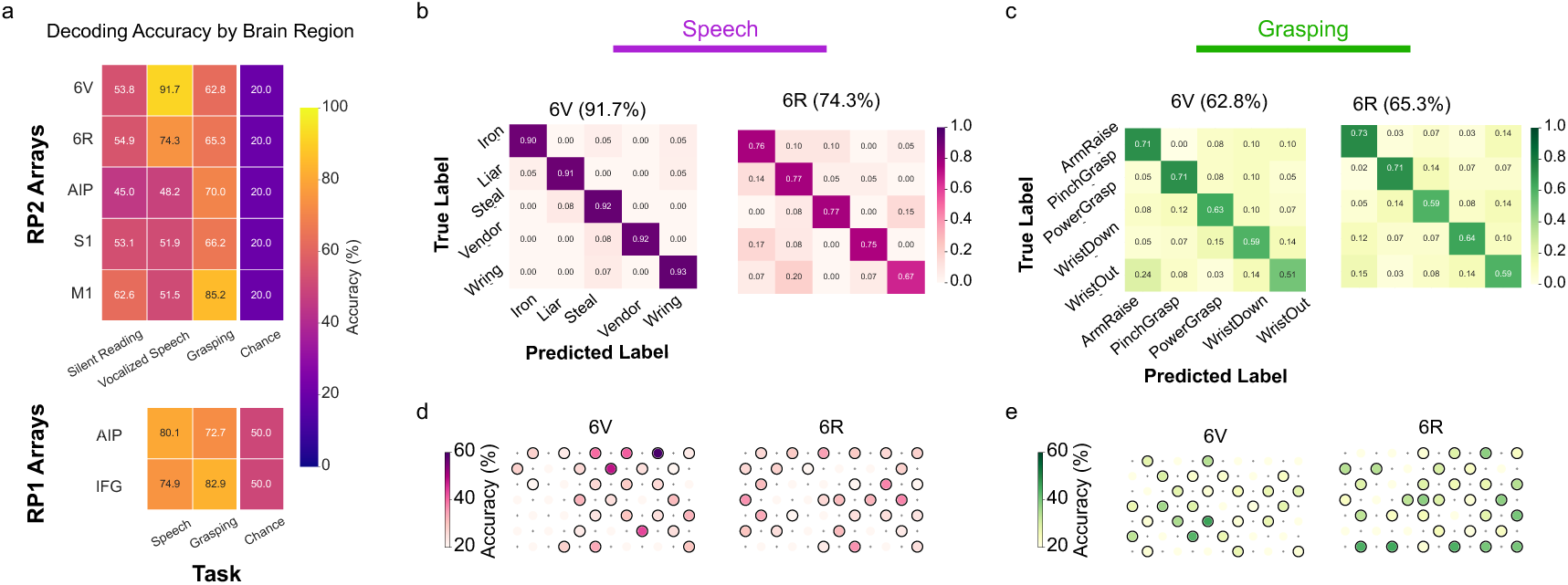
Neural decoding of speech and grasping. **a)** Task decoding accuracies based on 10-fold cross validation. Support vector machine (SVM) classifiers were trained to predict task conditions for both speech and motor tasks based on mean spike band power features from an optimally selected electrode subset (see Methods). Speech decoding involved discrete-word decoding from neural activity during covert (i.e., silent reading) and overt (i.e., vocalized) speech. For participant RP1, binary grasp and speech decoding (overt vs. covert) were performed. **(b)** Confusion matrix based on population decoding of vocalized speech in 6v (left) and 6r (right). These correspond to the results in **(a).** The corresponding classification accuracies are enclosed in parentheses. Confusion matrices for silent speech decoding can be found in supplementary Fig. 4. **(c)** Same as **(b)**, for grasping. **(d)** Heatmap showing electrode-level decoding accuracies for vocalized speech in 6v (left) and 6r (right). Circled electrodes showed selective task representation (i.e., decoding accuracy>chance level. Chance level= 20% for both tasks). **(e)** Same as **(d)** for the grasping task.

Overall decoding accuracies for silent speech in PMv were slightly lower than for overt speech (Fig. 2a; also see confusion matrices for 6r and 6v in supplementary figure 4). All subsequent analyses comparing speech and grasping use neural activity during vocalized speech. In RP2, decoding performance revealed opposite task biases across regions: PMv areas (6r and 6v) showed significantly better representation of speech than grasping (mean accuracy difference, speech – grasping = +19.0% ± 10.0 standard deviation, across arrays) whereas AIP, S1 and M1 favored grasping ( speech – grasp = −23.3% ± 8.0; Fig. 2a). We also quantified how well both speech and grasping are represented at the electrode level by running the decoding pipeline outlined above using features from each electrode separately. We then quantified the number of electrodes in each area with selective task representation (i.e., decoding accuracy > 20% chance level). Tuning heatmaps showing task encoding at the electrode level in 6r and 6v are shown in Fig. 2 (d, e). A significant fraction of electrodes across arrays showed above-chance representation of speech and grasping (for example, 58.1% and 76.7% of 6r electrodes as well as 64.3% and 57.1% of 6v electrodes showed better-than-chance tuning to speech and grasping, respectively.

Together, these results indicate that neural populations across motor, premotor, somatosensory and parietal cortices encode task-relevant information for both speech and grasping with high fidelity.

### Within-region functional connectivity reconfigures between speech and grasping

Having established that both speech and grasping are robustly represented across areas, we sought to quantify the functional structure of the population between tasks to better understand how both tasks are encoded. Prior work shows that the same neural population can markedly reconfigure its functional structure between behavioral states (e.g., movement planning vs. execution) (27). We asked whether the pattern of interactions among electrodes (i.e., coupling) is preserved or reorganized between tasks to support distinct computations. If coupling were task-invariant, electrode pairs that co-fluctuate during one task should exhibit comparable coupling during the other; conversely, distinct interaction patterns between tasks would imply a reconfigured functional architecture.

To test this, for each cortical area and task (speech, grasp), we computed the pairwise cross-condition correlations between electrodes (see methods) (Fig. 3a,b). This captures the functional structure (i.e., within-region functional connectivity pattern) for each task separately, revealing which electrode pairs behave similarly within each task. If neural populations have a fixed, task-invariant functional connectivity structure, the pairwise relationships between electrodes should remain largely unchanged between tasks. In this scenario, the correlation structure observed during one task (e.g., motor execution, Fig. 3a) would strongly predict the correlation structure during the other task (e.g., language processing, Fig. 3b), suggesting strongly overlapping neural subspaces. Visual assessment of pairs of correlation matrices (i.e., speech versus grasping) across multiple arrays reveal stark dissimilarity (Fig. 3 a, b).

**Figure 3.**
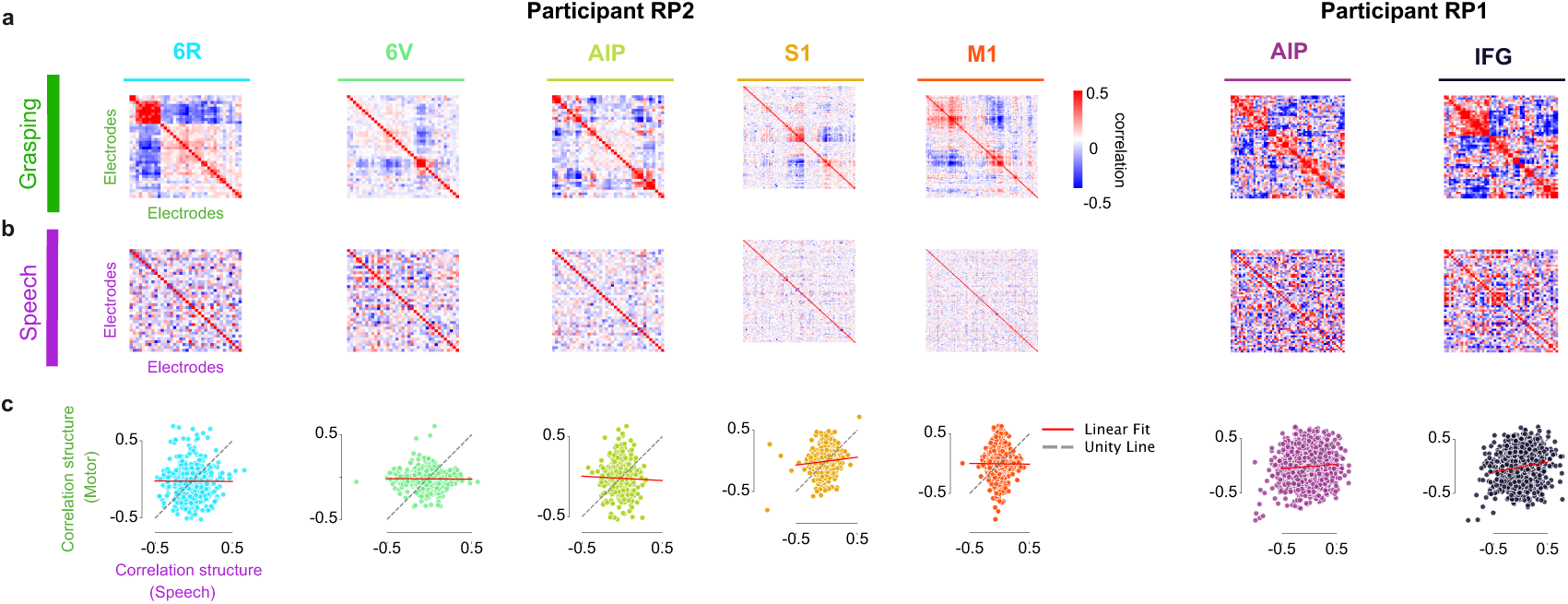
Cross-task reconfiguration of neural correlations. **(a, b)** Cross-condition correlations between pairs of electrodes within each array during speech **(b)** and grasping **(a)** tasks. Electrode order is identical between speech and grasp correlation heatmaps, highlighting the marked dissimilarity in within-area functional connectivity patterns between tasks. Only significantly tuned electrodes were included (i.e., electrodes with better-than-chance decoding accuracy in at least one task). **(c)** Scatter plot of vectorized (i.e., upper triangle of) correlation matrices in **(a, b)**. Linear regression was used to fit the grasp correlation structure as a function of the speech correlation structure (red).

To quantify this cross-task dissimilarity, we plotted the vectorized correlation matrices (upper triangle, excluding diagonal) between tasks against each other (Fig. 3c). We also applied linear regression to predict the correlation structure during grasping from the correlation structure during speech (Fig. 3c, red traces). If the pattern of correlations between electrode pairs remains identical between tasks (even if neural activity changes), data points would lie along (or close to) a unity line (Fig. 3c, dashed gray lines). Our results revealed negligible similarity between task-specific correlation structures: the regression of pairwise correlations between speech and grasping showed a near-zero fit (mean correlation coefficient = 0.011 ± 0.076 standard deviation, across all arrays; Fig. 3c).

These findings suggest that neural populations across multiple brain regions exhibit remarkable flexibility, allowing for task-dependent reconfiguration of within-region functional connectivity patterns to support new task demands.

### Co-representation of speech and grasping is facilitated by mixed selective neural populations

Next, we asked if the same neural population within each area contributed to both tasks, or whether different neural subsets are preferentially recruited between tasks. It is possible that the observed shift in population correlation structure is due to the recruitment of distinct neural subsets between tasks.

To investigate this, we quantified how “*active*” each electrode is within each task by computing its task variance (TV), which captures the strength of condition-dependent modulation of neural activity (see methods). A high TV value for a given task is indicative of statistically reliable, selective tuning across conditions of the task (e.g., different grasp types), whereas a lower value suggests weak, condition-independent response to task conditions. Plotting speech TV against grasp TV provided a population-level view of tuning: electrodes lying on or near the unity line exhibit comparable selectivity across tasks (Fig. 4b), whereas points below the line and toward the bottom-right indicate grasp preference, and points above the line and toward the top-left indicate speech preference. In 6v, most electrodes lay above the diagonal, indicating stronger modulation by speech than grasp. In all other brain regions, points were more dispersed, consistent with greater heterogeneity in task preference (Fig. 4b).

**Figure 4.**
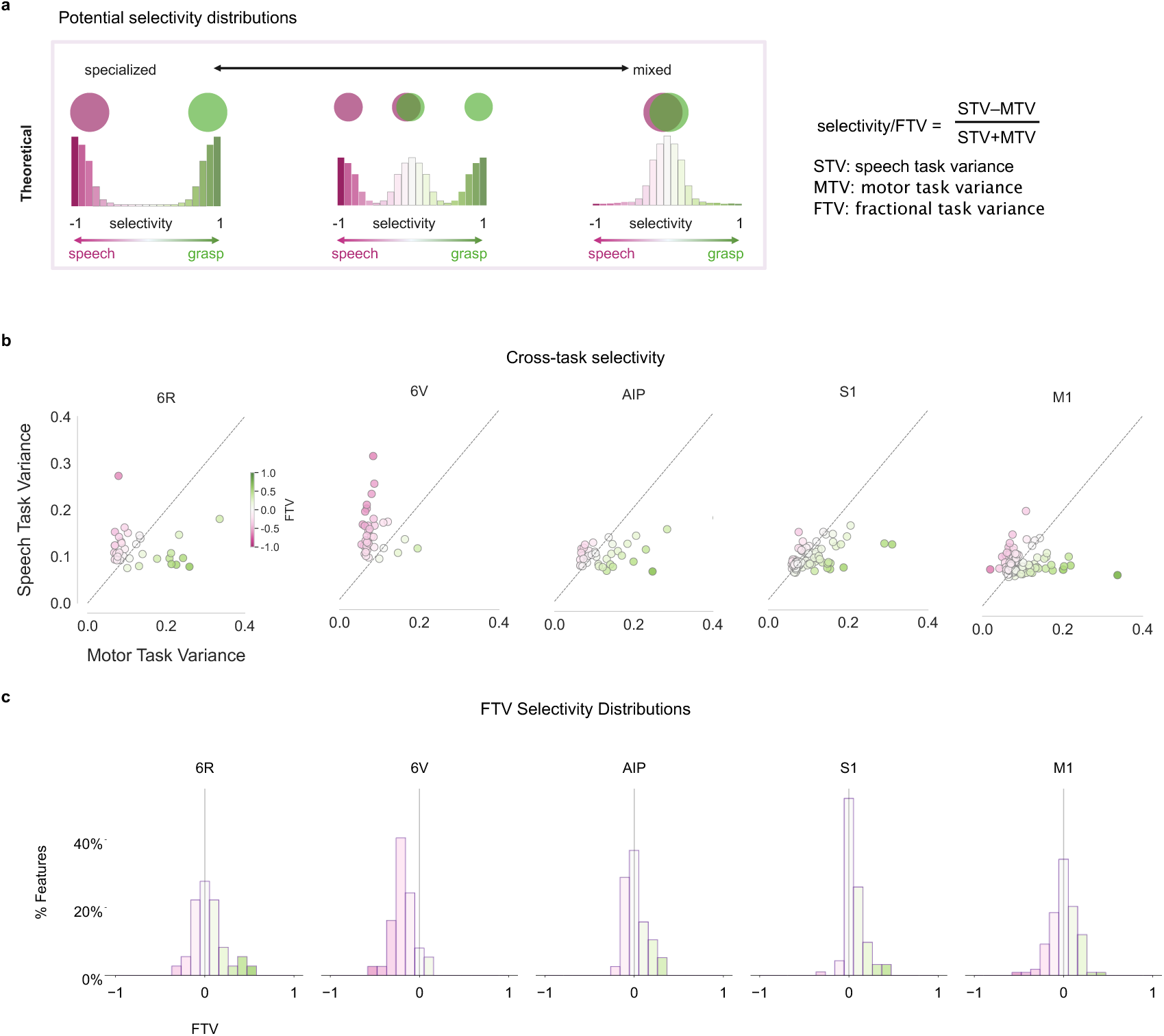
Relative selectivity for speech and grasping selectivity. **(a)** Schematic illustrating expected population-level selectivity distributions under different coding schemes, as assessed using Fractional Task Variance (FTV) analysis. For each electrode, FTV quantifies the relative difference in modulation strength (i.e., task variance, TV) between speech and grasping, normalized by total modulation (sum of task variances). FTV values range from −1 to +1, where −1 (under our convention) indicates preferential speech selectivity, +1 indicates preferential grasp selectivity, and 0 reflects mixed selectivity. For the entire array, if speech and grasping are encoded by non-overlapping, task-specific subpopulations, FTV yields a bimodal distribution (left). A unimodal distribution centered near zero indicates a mixed-selective population (right), while a multimodal distribution suggests a hybrid coding scheme (middle). Overlapping circles depict a single population showing mixed selectivity to both tasks. **(b)**Task variance scatter plots across RP2 arrays. Each dot represents a single electrode. The horizontal axis denotes TV during grasping, and the vertical axis denotes TV during speech. The unity line marks equal modulation across tasks; points above indicate stronger speech tuning, and points below indicate stronger grasp tuning. Colors reflect task selectivity (FTV). Only significantly tuned electrodes (i.e., those exceeding chance-level decoding accuracy in at least one task) were included. **(c)**Empirical FTV distributions across arrays. Each distribution pools electrodes within an array, providing an area-specific summary of relative task preference

To formalize each electrode’s relative involvement between tasks, we quantified cross-task selectivity using the Fractional Task Variance (FTV) metric introduced by Yang et al (28), which leverages TV measures between tasks to assign each electrode an integer value between 1 and –1 indicating task preference. Under our convention, an FTV score of 1 (or –1) denotes exclusive selectivity to grasping (or speech), and 0 denotes equal selectivity to both tasks. The task variance plots in Fig 4b were colored by electrode-level FTV scores to highlight tuning preferences of individual electrodes across different arrays. Since FTV values near zero could either indicate equally strong or equally poor tuning between speech and grasping, we identified and excluded electrodes that were not tuned to either task (i.e., decoding accuracy < chance level). We computed the FTV values for all electrodes and pooled the results into an FTV distribution highlighting the predominant tuning behavior of each brain region between the two tasks.

We considered three coding schemes by which the same neural population could represent two tasks (Fig. 4a): (i) *Functional specialization*, where distinct neural subpopulations are recruited for each task. This mechanism would yield a bimodal FTV distribution with two prominent peaks at the extremes corresponding to the two distinct classes within the neural population (Fig. 4a, left). (ii) *Mixed selectivity*, where both tasks engage largely the same neural population, predicting a unimodal FTV distribution (Fig. 4a, right). (iii) A *hybrid model*, where functional specialization and mixed selectivity coexist, yielding a multimodal FTV distribution (Fig. 4a, middle). It is worth noting that selectivity distributions derived from FTV analysis reflect an aggregate measure of task preference obtained by pooling single-channel selectivity values. As such, they are not necessarily predictive of population-level decoding performance. For instance, a population exhibiting equal task variance across two tasks (but characterized by redundant neural variance in one task and synergistic interactions in the other) could yield a zero-centered unimodal selectivity distribution while producing markedly different decoding outcomes between tasks.

Our results revealed unimodal FTV distributions in all areas (Hartigan’s dip test for unimodality: 6r, p = 0.83; 6v, p = 0.53; AIP, p = 0.23; S1, p = 0.96; M1, p = 0.84) (Fig. 4c), arguing against segregated subpopulations and in favor of mixed selectivity. The means of the selectivity (i.e., FTV) distributions in 6v, S1 and AIP did not center at zero (one-sample t-test, 6v, p<0.01; S1, p<0.05; AIP, p<0.05), indicating a bias toward speech in 6v and a slight bias toward grasping in AIP and S1 under the stated sign convention. The distribution means in 6r and M1 did not significantly differ from zero (one-sample t-test: 6r, p = 0.94; M1, p = 0.67), indicating a more balanced involvement of the full neural population across tasks (Fig. 4c).

Together, these findings support a model where speech and grasping are co-represented by a predominantly overlapping, mixed-selective population in each region, with modest area-specific biases.

### Mixed selective neural populations exhibit structured spatial organization

Our results so far are consistent with the idea that cortical networks largely contribute to both tasks rather than forming two segregated, task-specific subpopulations. However, task selectivity distributions only indicate *which* electrodes contribute and to what extent, not *where* those contributions concentrate spatially on the array. Mixed selectivity at the population level can still coexist with subtle topographic biases; for example, weak spatial gradients or small clusters that a global selectivity histogram would miss. To address this, we selected three arrays showing different biases in selectivity distributions (AIP, with a slight grasp preference; 6v showing speech bias and 6r with a more balanced selectivity distribution) and asked whether the observed task biases have a spatial signature on the arrays. For example, do the most active electrodes for each task (i) cluster spatially into an “information hotspot”? (ii) If so, are these information hotspots spatially distinct between tasks?

To accomplish this, for each task, we identified the most task-informative electrodes (prediction accuracy > 25%) and computed a weighted spatial centroid using leave-one-out cross-validated electrode accuracies as weights while considering each electrode’s 2D coordinates on the array. We overlaid these centroids on the 7×12 array grid (Fig. 5a,c,e), to reveal the spatial distribution of information hotspots between speech and grasping across all arrays. For a one-dimensional summary, we projected points onto a linear-discriminant analysis (LDA) task-separating axis (Fig. 5b,d,f).

**Figure 5.**
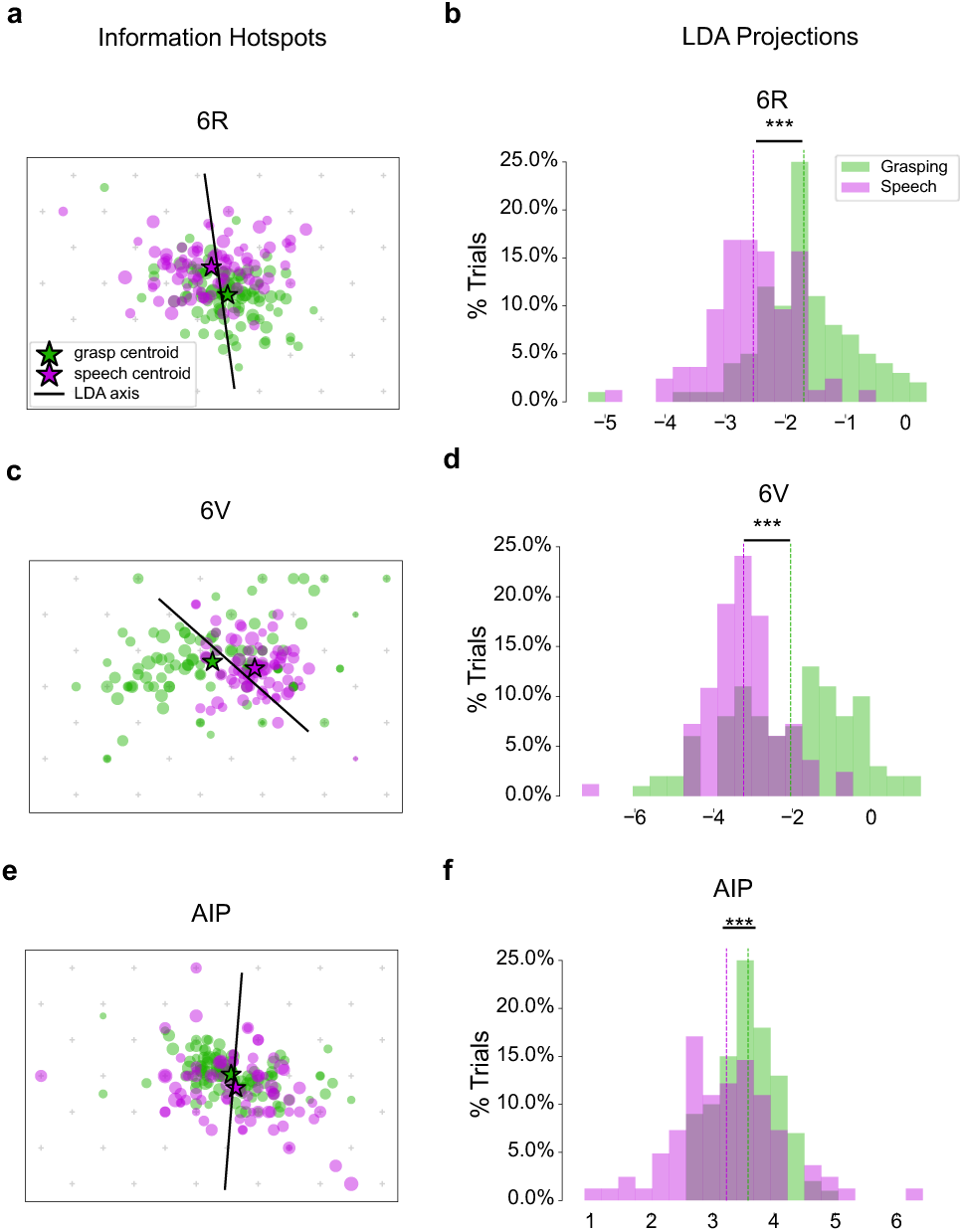
Spatial organization of speech and grasp representations. **(a)** Task-specific information “hotspots” overlaid on 6r array map. Gray markers (**+**) indicate electrode coordinates. Information hotspots were estimated by computing a weighted centroid from the individual electrode decoding accuracies, taking the 2D electrode coordinates into consideration. Each dot shows the centroid computed from one cross-validation fold for each task. LDA was used to identify a task-separating axis (black line) within this scatter plot distribution of weighted centroids. **(b)** Projections of the cross-validated weighted centroids onto the LDA axis for 6r. **(c–f)** Same as **(a, b)** for 6v and AIP arrays in RP2.

Despite the unimodal FTV distributions (consistent with mixed selectivity), all arrays showed reliable spatial offsets between speech and grasp centroids as assessed via permutation testing (p < 0.001) (Fig. 5d) This spatial separation of information hotspots between tasks is compatible with a dual task encoding model that combines mixed selectivity with a mild topographical organization of neural populations rather than strict intermixing or hard segregation.

In other words, both tasks generally draw on a common neural population yet task-informative neural activity peaks in different spatial zones for speech versus grasping.

### Neural representations of speech and grasping occupy distinct, non-orthogonal subspaces

Building on the observed cross-task reconfiguration of pairwise correlations, we asked whether language and motor computations occur in compartmentalized (i.e., task-specific) neural subspaces or in a shared subspace. In contrast to pairwise and single-channel metrics, subspace analysis characterizes the low-dimensional geometry of population activity, revealing possible task-specific rotation or segregation of neural manifolds even when individual channels exhibit mixed-selectivity. For each area and task, we applied principal component analysis (PCA) to the cross-condition neural activity to define a ten-dimensional task subspace (i.e., top 10 eigenvectors of the task-specific covariance matrix; see Methods). We first quantified within-task capture by projecting each task’s neural activity onto its own subspace and computing the fraction of total variance explained (Fig. 6). This provided a summary of how well the full-dimensional neural data for each task is represented by its corresponding low-dimensional task subspace. Example projections of 6r speech activity onto the top 3 principal components (PCs) within the speech subspace are shown in Fig. 6a (top row). Similar projections of grasp activity onto the grasp subspace are shown in Fig. 6b (bottom row). We also quantified cross-task capture by projecting activity from one task onto the other task’s subspace to assess how well the two geometries align. Example projections of grasp activity onto the speech subspace are shown in Fig. 6a (bottom row). Similar projections of speech activity onto the grasp subspace are shown in Fig. 6b (top row).

**Figure 6.**
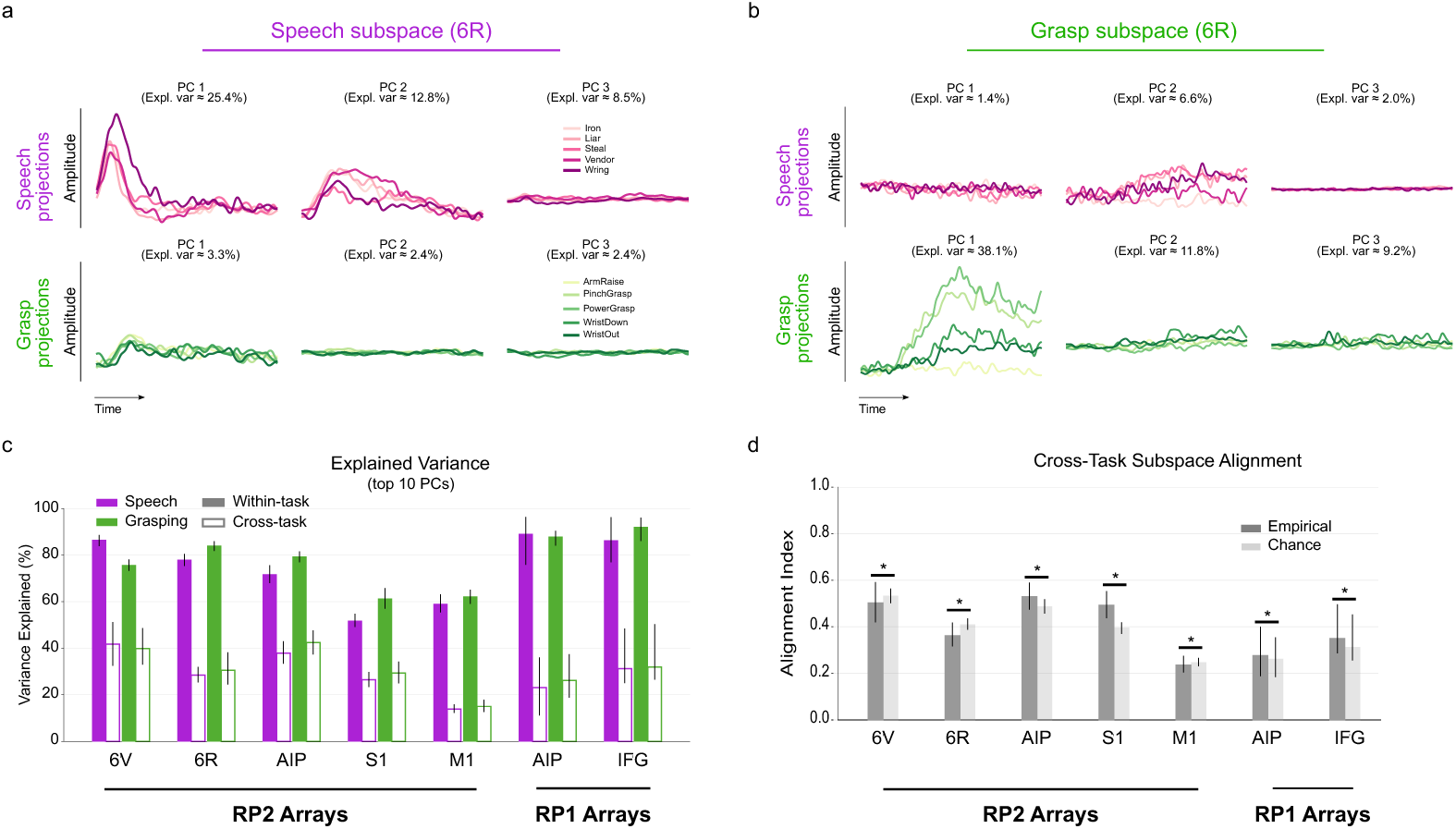
Speech and grasp subspace segregation. **(a)** Projections of speech and grasp neural activity onto speech subspace in 6r. PCA was used to derive a 10-dimensional speech subspace. For visualization, projections of speech (top) and grasp (bottom) activity onto the top three PCs are shown, together with the variance captured by these projections. **(b)** Projections of speech (top) and grasp (bottom) neural activity onto the grasp subspace in 6r. **(c)** Within- versus cross-task cumulative variance capture. Bars reflect the cumulative variance captured when projecting neural activity onto task-derived 10-dimensional subspaces. “Within-task” refers to projecting each task’s activity onto its own subspace; “cross-task” refers to projecting one task’s activity onto the other task’s subspace. **(d)** Cross-task subspace alignment. Alignment indexes quantify cross-task cumulative variance captured as a fraction of within-task cumulative variance (0–1 scale; higher indicates greater overlap). Empirical alignment values are compared against a null distribution obtained by projecting onto randomly sampled subspaces within the same neural state space (significance assessed via permutation). An alternative view of these results is provided in supplementary Fig. 6.

Across all brain regions, within-task subspaces captured a large fraction of their own task variance (i.e., cumulative variance across top 10 PCs) in both speech and grasping tasks (Fig. 6c, solid bars). In contrast, cross-projecting grasp activity onto speech subspaces (or speech activity onto grasp subspaces) resulted in substantially less captured variance (Fig. 6c, hollow bars), suggesting poor generalization of subspaces between tasks. In other words, the dominant axes for one task are not well explained by the other task’s subspace. The degree of overlap (or lack thereof) between speech and grasp subspaces was directly computed using a subspace alignment index (27) that expresses the cross-projected variance capture as a fraction of within-task variance capture (0–1 scale; higher = more overlap). Empirical values were compared against a chance baseline obtained from projecting neural activity onto randomly oriented subspaces in the same geometric space (see Methods).

Statistically, empirical alignment was slightly lower than baseline (left-sided paired t-test, p<0.05) in 6v, 6r and M1, and slightly higher than baseline (right-sided paired t-test, p<0.05) in S1, IFG and AIP (both participants). Nevertheless, the absolute deviation from baseline was close to zero across arrays (mean absolute deviation, |empirical–baseline| = 0.042 ± 0.026, across all arrays; Fig. 6d, supplementary Fig. 6). This small absolute difference between empirical and chance alignment indicates that speech and grasping occupy largely distinct subspaces within each area (i.e., subspaces with minimal overlap comparable to chance levels).

## Discussion

This study highlights the remarkable dual functionality of multiple cortical regions, demonstrating the capacity of motor, premotor, frontal, somatosensory, and parietal cortices to encode both motor and language functions. The functions of these regions have often been framed within the context of motor-planning, execution and somatosensory feedback but our results align with growing evidence that cortical representations of multiple functions (particularly speech and motor intentions) overlap (11, 13, 14, 16, 29).

Our findings within PMv reveal that while both subregions exhibit an intermixed representation of speech and grasping, 6v exhibited more speech dominance relative to grasping. These observations are consistent with previous findings demonstrating that while 6v is surprisingly broadly tuned to whole body movements, at a finer scale, it is most selective to speech (16). On the other hand, our findings in 6r address a significant knowledge gap in its function in the human cortex, revealing its dual functionality in both speech and motor processing.

Theoretically and computationally, mechanisms for multitask representation have been proposed to lie along a specialization–flexibility continuum (28, 30) (Fig. 4c). On one end of this spectrum, functional specialization drives the formation of network modules, which are groups of highly specialized neural clusters that compute specific task variables. In this view, the integration of computations across specialized modules allows a single network of neurons to efficiently represent multiple task parameters. Anatomically, one might expect spatially segregated clusters for speech versus hand control in certain areas, especially given recent evidence suggesting the ventral region of the human premotor cortex could be subdivided into four areas (6v1, 6v2, 6v3, 6r) based on cytoarchitectonic similarity (31). Within this framework of functional segregation, one could expect minimal functional (and possibly, spatial) overlap between the distinct neural populations activated by different tasks.

On the flexibility end of the spectrum, mixed-selective representations emerge at the level of individual neural elements (e.g., single channels) within the network. In this framework, cortical neural populations across species have been found to exhibit a remarkable versatility that allows them to encode information about multiple tasks (32, 33) and different parameters of the same task (17, 34–36) with high fidelity. An immediate benefit of this organization lies in the flexibility it confers, allowing the brain to economically implement multiple readouts of the same neural population to support different task demands.

Our task selectivity analyses place all areas nearer the flexibility end of this spectrum. In each area, the task selectivity distribution (based on FTV) was unimodal, arguing against segregated, task-specific subpopulations. This implies most neural elements in all areas contribute to both tasks, with modest biases. For BMIs, this means overlapping electrode sets can carry both motor and speech information, improving adaptability and efficiency. This has a potential for minimizing the number of distinct cortical implants needed in patient applications requiring joint decoding of both tasks (as in the case of patients with locked-in syndrome).

The observed mixed selectivity was driven by the ability of neural populations to markedly reconfigure their *“population codes”* (i.e., correlation structures) between tasks. Indeed, we observed task-dependent reconfiguration of functional coupling, with pairwise correlations differing substantially between tasks, consistent with transient relaxation of network constraints to meet new computational demands (37). This ability to reorganize functional architecture is consistent with previous studies in large-scale functional brain networks such as the fronto-parietal cognitive control network (FPN), and underscores the brain’s capacity to optimize its networks for diverse tasks, as demonstrated by multiple studies (38–40). Together, these studies reveal that multi-task representation across distributed cortical networks is driven primarily by flexible hubs, brain regions (primarily in the FPN), exhibiting variability in task-dependent connectivity with multiple specialized networks (such as auditory and visual networks). Our findings here suggest that localized brain regions (such as PMv) capable of multi-task representation, on a smaller scale, might be able to do so by mimicking the flexible functional reconfiguration observed in distributed, adaptive networks such as the FPN.

Mixed selectivity at the population level, however, does not preclude structured spatial or anatomical organization based on distinct tuning properties. In our data, the most informative channels for speech and grasping showed small but reliable spatial offsets on each array. This is not inconsistent with mixed selectivity; rather, it suggests a graded topography (i.e., overlapping populations with spatial shifts in peak information density) instead of distinct neural modules for speech versus grasping. Conceptually, this pattern is plausible: PMv and neighboring regions exhibit known somatotopic and articulatory gradients (13, 16), and prior work has reported patchy, partially overlapping maps for hand/face and speech-related effectors as well as gradients in tuning to whole-body movements along the precentral gyrus across multiple individuals (16).

For BMI applications, the spatial biases reported here are actionable. First, they support task-aware channel selection: emphasizing speech-leaning patches for word decoding and grasp-leaning patches for motor control can boost performance. Second, they motivate spatial priors in multi-functional decoders (e.g., mild L1/L2 penalties that respect array topology), improving generalization and robustness to noise. Third, the offsets argue for adaptive, state-dependent weighting, allowing the same implant to reweight channels when switching between speech and grasping, rather than training wholly separate models or requiring distinct implants in multiple cortical regions.

Geometrically, subspace analyses revealed distinct but non-orthogonal manifolds for speech and grasp computations within each area. Intrinsic (i.e., within-task) projections captured substantially more variance than extrinsic (i.e., cross-task) projections, indicating compartmentalized computations for speech and grasping. These findings align with previous work demonstrating compartmentalized computations in both dorsal premotor (PMd) and primary motor (M1) areas in non-human primates (27, 41, 42). Movement preparation and execution were shown to occupy near-orthogonal subspaces in both PMd and M1 (27), and similar separations have been observed in PMd when considering the representations of multiple movement types (42). Additionally, the neural representations of action observation (covert movement) and movement execution have been shown to exhibit a significant overlap that could be separated out into mutually orthogonal subspaces through a joint manifold optimization (41), reinforcing the view that task demands modulate population geometry rather than recruiting entirely distinct neural populations. Taken together, these results support subspace segregation as the primary mechanism by which mixed-selective neural populations minimize interference between computations.

## Materials and Methods

### Study participants

Two participants with chronic cervical SCI were enrolled in the ReHAB clinical trial (ClinicalTrials.gov ID: NCT03898804), conducted under an Investigational Device Exemption from the U.S. Food and Drug Administration and approved by the local Institutional Review Board. Informed consent was obtained from both participants prior to enrollment. Participant RP1 was a 27-year-old right-handed male at the time of implantation with a C3–C4 level American Spinal Injury Association Impairment Scale B (AIS-B) SCI. Participant RP2, a 47-year-old right-handed male, sustained a C3–C5 AIS-B SCI. As part of the trial, each participant received intracortical microelectrode implants targeting sensorimotor and association cortical regions involved in hand and arm function. In RP1, microelectrode arrays were implanted in the “hand knob” area of the primary motor cortex (M1) and primary somatosensory cortex (S1), as well as in the anterior intraparietal area (AIP) and inferior frontal gyrus (IFG).

Each of the AIP and IFG sites were each implanted with a single 8 × 8 (64-channel) 1.5-mm iridium oxide microelectrode array (Blackrock Neurotech, Salt Lake City, UT). We targeted the medial junction of the intraparietal and postcentral sulci (18, 43–45) for AIP, and the anterior bank of the border of area 44 and ventral premotor cortex for IFG. AIP implantations were guided by well-established anatomical landmarks (18, 43, 45), while the IFG site was functionally localized using intraoperative awake speech mapping to avoid language-critical areas (18). RP2 received a total of eight arrays spanning five cortical regions. This included three arrays in S1 (a total of 128 electrodes in a 43/42/43 split), two arrays in the hand knob of M1 (128 electrodes in 64/64 channel splits), and three additional arrays targeting AIP, 6v, and 6r (128 electrodes in a 43/42/43 split, respectively). Surgical targeting of PMv areas leveraged a multi-modal approach combining MRI-based anatomical mapping, task-based functional imaging via fMRI, and 3D surgical modeling (46).

### Task design

Experiments were conducted in 3–4-hour sessions composed of multiple blocks (2–10 minutes each). Participants remained seated upright in their wheelchairs, facing a monitor that displayed visual cues and instructions for each trial.

#### Motor Tasks

For RP1, each trial began with the display of a semi-transparent “ghost hand” for 1.5–2 seconds, followed by a “go” phase during which a video of a solid “control hand” performed an opening or closing movement (see supplementary Fig. 1, bottom). RP1 was instructed to imagine executing the depicted hand movement. These open-loop trials were used to train a Kalman filter for one-degree-of-freedom aperture decoding. In subsequent closed-loop blocks, the trained decoder allowed RP1 to control the grasp aperture of the control hand in real time. Only data from closed-loop trials were used in analyses.

For RP2, trials began with a 3–3.5 second presentation of a text cue describing a specific arm or hand movement to attempt (e.g., wrist extension, hand squeeze). A go cue then prompted RP2 to initiate the instructed movement, followed 3 seconds later by a return cue signaling the participant to relax to a resting position (Fig. 1b). Movement types were randomized across trials to mitigate anticipatory effects.

#### Language Tasks

In language trials, both participants engaged in tasks with defined covert and overt phases. RP1 was presented with a phrase cue consisting of a sentence with a blanked-out verb, followed by a word cue showing the base form of the missing verb. The participant was instructed to covertly conjugate the verb to match the grammatical context of the phrase. A color change then served as a go cue, prompting RP1 to overtly articulate the conjugated verb. The phrase and word cue epochs defined the covert phase, while the overt phase corresponded to the verbal response following the go cue (see supplementary Fig. 1, top).

For RP2, each trial began with a 1.5-second rest epoch, followed by a covert instruction epoch lasting 1.5–2 seconds, during which a single word appeared on the screen. RP2 was instructed to silently read and retain the word in memory. Immediately following this, a go cue prompted the participant to overtly speak the word. Word identity was randomized across trials to reduce predictability (Fig. 1b).

### Extraction of neural features

Broadband neural signals were recorded at 30 kHz from the microelectrodes in all implanted areas using the Neuroport system (Blackrock Neurotech, Salt Lake City, UT) while subjects performed the motor and language tasks. Offline preprocessing consisted of bandpass filtering the raw signals between 0.25 and 5 kHz with a zero-phase 4th-order Butterworth filter. Spike Band Power (SBP) features were extracted by computing the mean squared amplitude of the filtered signal within non-overlapping 10ms bins. To mitigate common-mode noise across electrodes, we applied spatial filtering using linear regression referencing. SBP features were smoothed with a Gaussian kernel (standard deviation = 30–50ms) and mean-centered within each block on a per-electrode basis. These processed features were used in all subsequent analyses.

### Neural decoding of speech and grasping

Decoding of task conditions was achieved using either a standard support vector machine (SVM, with a radial basis function kernel) or an ensemble stacking model consisting of two base learners—an SVM and a Gaussian naïve bayes classifier—combined via a logistic regression meta-classifier. Only speech classification in RP1 used the ensemble model. Classification was performed on mean neural activity within task-specific time windows. In RP2, this window was 2s prior to the go cue for grasping, and 1.5s following the go cue for speech, offset by 200ms to account for reaction time. In RP1, it was 1s after go-cue during grasping and 1.5s before and after speech onset cue during the speech task.

To reduce dimensionality and maximize decoding performance, we employed a sequential forward selection strategy (47) (see supplementary Fig. 3). First, the electrode with the highest individual decoding accuracy was selected. Additional electrodes were then iteratively added to minimize prediction loss, with the procedure terminating once the maximum number of electrodes is reached or when no improvement was observed after 10 iterations. This approach yielded a compact feature set composed of the most informative and synergistic subset of electrodes. Model performance was assessed using either leave-one-out cross-validation or 10-fold cross-validation and compared against a baseline chance level accuracy of 1/C, where C is the number of unique task conditions.

### Pairwise electrode correlation analysis

To assess the functional organization of intracortical activity during different task contexts, we computed pairwise correlations between electrodes within each array and task condition using a data structure similar to that described by Elsayed et. al (27). For grasping, windowed neural features were organized in the matrix 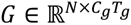, where 𝑁 is the total number of electrodes, 𝐶_g_ is the number of unique grasp types, and 𝑇_s_ is the total number of bins within the selected grasp window. Thus, this procedure averages the time-series features across trials for each grasp type separately and unrolls the data across all conditions into the 2D matrix G. This procedure was applied to each array separately. For the speech task, an analogous matrix 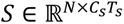 was constructed where 𝐶_s_is the number of unique speech conditions (unique words), and 𝑇_s_ is the total number of bins within the selected speech window. For both tasks, Pearson’s correlation coefficients were computed between each pair of electrodes (i.e., rows of 𝑆 or 𝐺) to generate an 𝑁 × 𝑁 correlation matrix per task and array. These matrices quantify the degree of shared tuning patterns across electrodes and serve as proxies for within-area functional connectivity.

To examine whether pairwise correlation structure was conserved or reconfigured across tasks, we compared the correlation matrices derived from the grasping and speech conditions. Specifically, the lower triangle of each correlation matrix (excluding the diagonal) was vectorized to form a feature vector for each task. Linear regression was then used to fit the speech-derived correlation vector as a function of the grasp-derived vector. The coefficient of determination and regression slope quantify the degree to which task-specific correlation patterns were aligned. A high correlation would suggest task-invariant functional coupling, whereas deviations from this relationship indicate context-dependent reorganization of local network structure. This analysis was repeated independently for each array and participant. The resulting regression lines are shown in Fig. 3c for all arrays.

### Task selectivity analysis

We quantified electrode selectivity across tasks using the Fractional Task Variance (FTV) metric introduced by Yang et al (28). This method characterizes the degree to which an electrode’s activity is preferentially modulated by one task relative to another. First, we computed the task variance (𝑇𝑉), which captures the strength of condition-dependent modulation of electrode activity within a given task. For electrode 𝑖 during the speech task, task variance, *TV_s_^i^*, was defined as:

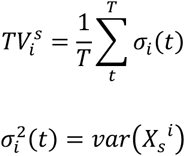

Where 𝑣𝑎𝑟 denotes the variance operator applied across trial-averaged conditions at each time point, 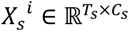 is the matrix of neural activity for electrode 𝑖 over the 𝑇_s_ speech window time points and 𝐶_s_ unique speech conditions. The variable 𝜎^2^(𝑡) stores the computed variance across time within the task window, whereas 𝜎(𝑡) is the standard deviation. A similar calculation yielded *TV_i_^g^* for the grasping task. From this, the preferential selectivity of electrode 𝑖 between speech and grasping was computed as follows:

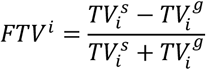

By construction, FTV values range from −1 to 1. A score of −1 indicates exclusive selectivity to speech *TV_i_^s^* ≫ *TV_i_^g^* whereas a score of 1 indicates exclusive selectivity to grasping *TV_i_^g^* ≫ *TV_i_^s^*. A value of 0 reflects mixed tuning, with equal cross-condition variance across tasks. To assess task selectivity at the population level, FTV values were pooled across electrodes to generate array-specific distributions, summarizing the overall task preferences within each cortical region (Fig. 4c).

### Analysis of spatial segregation

To determine whether information about speech and grasping was spatially localized within the implanted arrays, we quantified electrode-wise decoding performance and identified task-related “information hotspots.” For each electrode, decoding accuracy was computed across leave-one-out cross-validation folds. These accuracies were then used as weights to calculate a spatially weighted centroid within each array, representing the locus of maximal task-relevant information while accounting for both electrode position and decoding strength.

Centroid locations were computed separately for speech and grasping tasks, and their relative displacement was measured. A significant separation between centroids would indicate spatial segregation of task-relevant representations, whereas overlapping centroids would suggest shared or intermixed encoding across the array.

To further characterize this organization, we applied linear discriminant analysis (LDA) to the centroid data. LDA projects data into a low-dimensional space that maximizes separation between task categories while minimizing within-task variability. By projecting centroid locations into this discriminant space, we obtained optimized representations of task-specific segregation. For visualization, projections onto the first LDA axis were plotted (Fig. 5a,c), providing an interpretable view of the degree of separation between speech- and grasp-related centroids.

### Dimensionality reduction and assessment of subspace overlap

We applied principal component analysis (PCA) to the electrode feature matrices for speech (𝑆) and grasping (𝐺) to obtain low-dimensional representations of neural activity. The full premovement epoch during grasping was used for the grasp PCA while a window of 2s after go cue (with a 200ms offset) was used for the speech PCA. For each task, we defined the neural subspace (or manifold) as the geometric space spanned by the top 10 principal components (PCs), denoted as 𝑆 − 𝑃𝐶𝑠 for speech and 𝐺 − 𝑃𝐶𝑠 for grasping. To evaluate how well each task’s neural activity was captured by its corresponding manifold, we projected speech data onto the 𝑆 − 𝑃𝐶𝑠 and grasping data onto the 𝐺 − 𝑃𝐶𝑠 and measured the proportion of variance explained. To quantify cross-task generalization, we then projected speech data onto the 𝐺 − 𝑃𝐶𝑠 and grasping data onto the 𝑆 − 𝑃𝐶𝑠, again measuring the variance captured. The similarity between the two task subspaces was quantified using the subspace alignment index (AI)(27). The alignment index measures how well the subspace derived from one task accounts for variance in the neural activity of the other task, relative to how well it explains variance in its own task. Based on this framework, the overlap of the motor subspace with speech subspace (a measure of how well motor subspace can explain the variance in speech activity) can be computed as follows

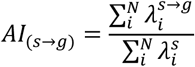

where 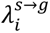 is the amount of variance in speech activity captured by projecting speech data onto the 𝑖^th^dimension in grasping subspace (𝑖^th^grasping PC), and *λ_i_^s^* is the variance in speech captured by its own 𝑖*^th^* PC. 𝑁 denotes the dimensionality of the subspace (which we set to 10). This can be reformulated as follows:

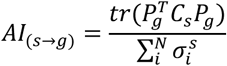

Where 𝑡𝑟 is the trace operator, 𝑃*_g_* is a matrix of the top 𝑁 grasping PCs (*P_g_^T^* being its transpose), 𝐶_s_ is the covariance matrix of speech features and 𝜎^s^ represents the singular values of 𝐶_s_. By construction, AI values range from 0 to 1, with higher values indicating stronger overlap between task subspaces whereas values close to zero indicate near-orthogonal subspaces. For interpretability, two subspaces are considered orthogonal if every vector in one subspace is perpendicular to every vector in the other (dot product = 0), corresponding to an AI value approaching zero.

### Estimation of baseline subspace alignment

Neural activity resides in a high-dimensional space, within which task-related neural manifolds represent constrained, low-dimensional subspaces. A key question is whether speech and grasping occupy distinct manifolds that exhibit strong misalignment, or whether apparent misalignment simply reflects the high dimensionality of neural activity, where randomly sampled subspaces are expected to be nearly orthogonal by chance. To test this, we generated random subspaces constrained by the covariance structure of the empirical data using the approach described in (27). Specifically, we projected the neural activity onto random sets of dimensions drawn from the same covariance structure as the speech (𝑆) and grasping (𝐺) feature matrices, thereby preserving pairwise covariance statistics while randomizing the orientation of the subspaces. We repeated this procedure over 1,000 iterations, each time computing the variance captured and the corresponding alignment index. The resulting null distribution provided a baseline expectation of subspace overlap given the dimensionality and covariance of the data. Specifically, the chance levels control for the size of the chosen subspace dimensionality (10, in this study) relative to the dimensionality of the full state space. Empirical AI values were then compared to this shuffled distribution to determine whether observed subspace overlaps significantly exceeded chance. Any empirical “overlap” that is comparable to chance could largely be attributed to the sizes of the task subspaces relative to the full state space dimensionality, as opposed any shared structure between the two subspaces.

## Acknowledgement

The authors wish to thank the study participants and their families for their time and commitment to this study. This research was supported by the following grants: VAMR 5I01RX002654, DoD CDMRP SCIRP SC180308.

---

**Supplementary Figure 1.**
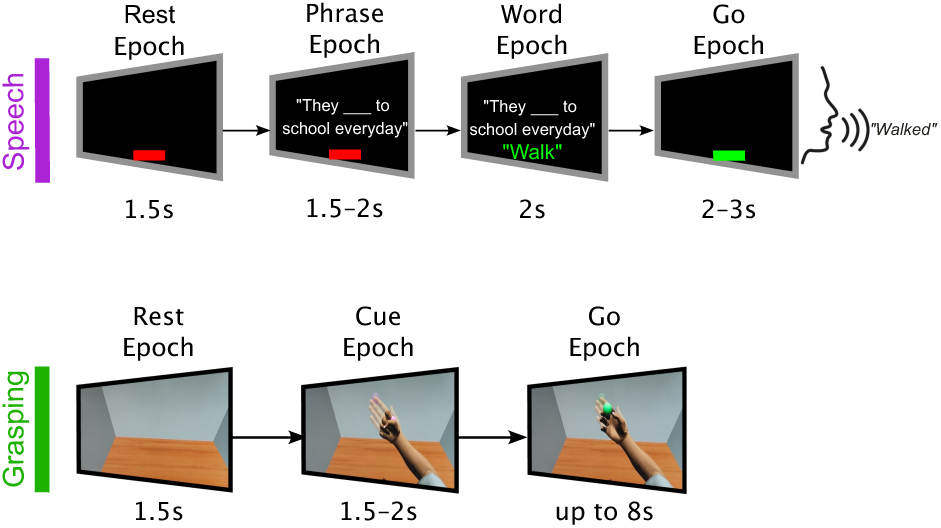
RP1 speech and grasp task design. During the speech task (top), RP1 performed grammatical inflection (i.e., verb conjugation) based on randomly cued phrase prompts. The task included (i) a covert phase, where the participant viewed a phrase with an omitted target verb and determined the appropriate conjugation, followed by (ii) a go phase, where they overtly articulated their conjugated response. The motor task (bottom) consisted of three phases: (i) a baseline phase with no movement, (ii) a cue phase where a visual cue displayed the target virtual grasp aperture (power grasp), and (iii) a go epoch where the participant performed 1-dimensional closed-loop control (opening or closing) of the virtual hand aperture

**Supplementary Figure 2.**
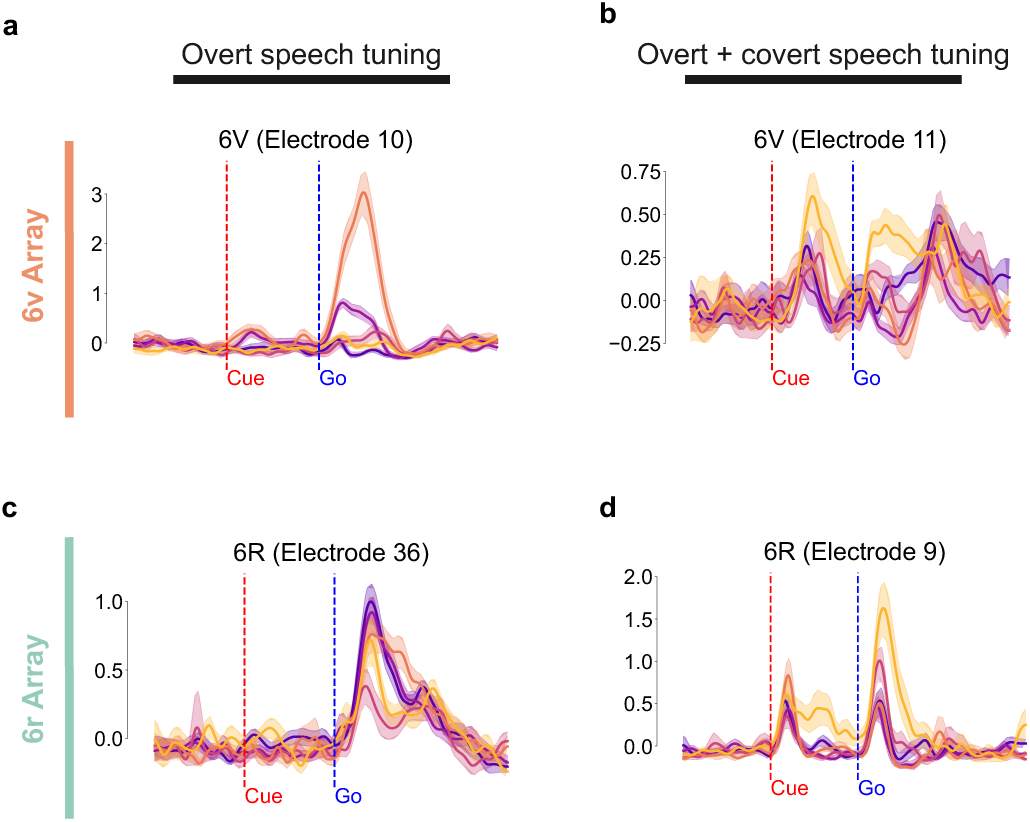
Additional example PMv features showing preferential tuning to overt (vocalized) and/or covert (silent) speech. Electrodes across areas showed a variety of rich tuning patterns during the speech task. For example, some electrodes (**a, c**) showed selective preference for speech vocalization (while remaining relatively inactive during silent reading). Other groups of electrodes (**b**, **d**) showed strong task-selective activity during both silent reading and vocalized speech.

**Supplementary Figure 3.**
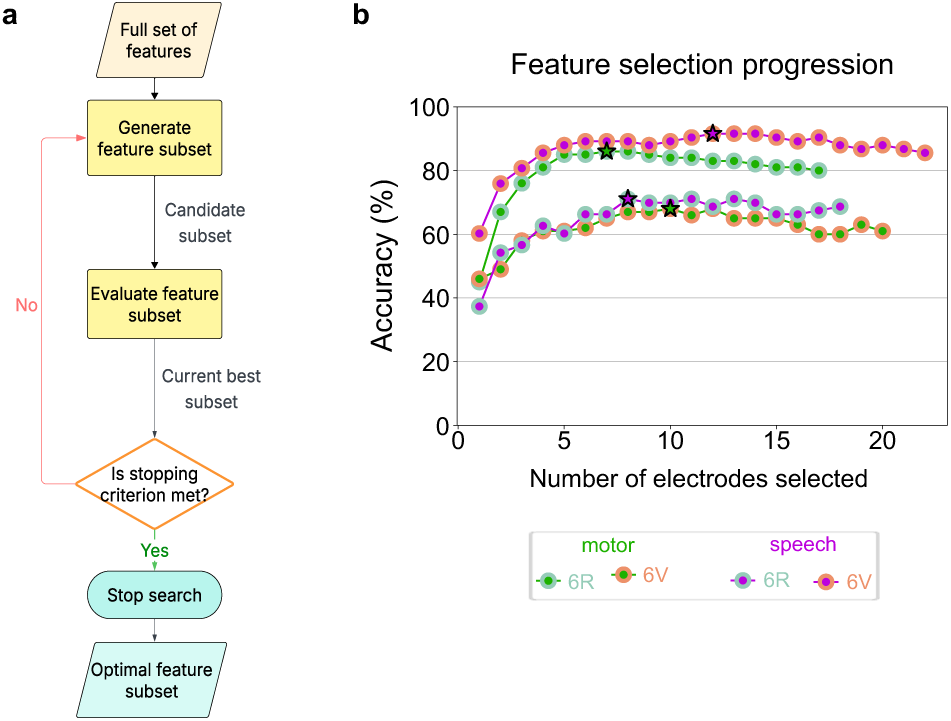
Sequential feature selection algorithm and example results. **(a)** Illustration of the procedure. We first evaluate single-feature decoding accuracy for every feature, rank features by accuracy, then greedily add features one at a time. Starting with the top-ranked feature, at each step we add the feature that yields the largest cross-validated gain (or least degradation) in accuracy when combined with the current set. The search stops when adding features fails to improve validation accuracy after ten iterations. The set of features yielding the best performance up to that point are returned as the optimal feature subset. **(b)** Example accuracy vs. number of features curve showing rapid gains with a small subset and saturation thereafter; the selected number of features based on cross-validation are highlighted for each array and task.

**Supplementary Figure 4.**
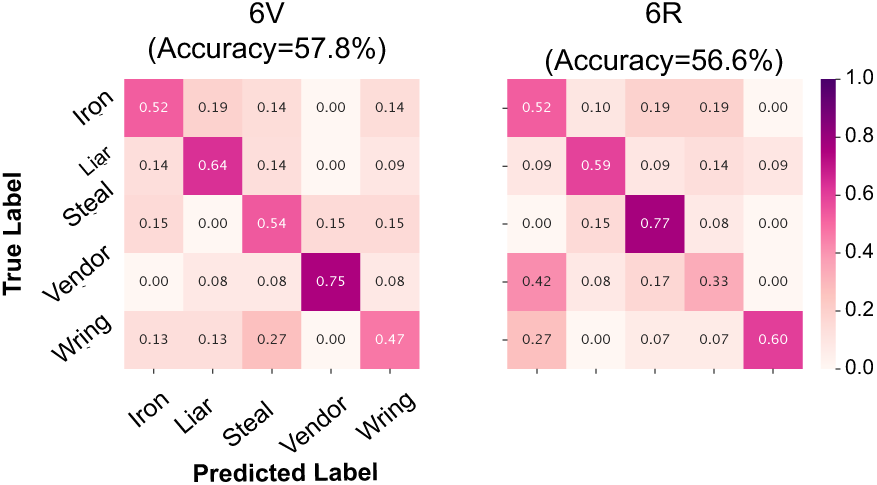
Confusion matrix from silent speech discrete-word decoding. Confusion matrices based on discrete-word speech decoding during silent reading across 6v (left) and 6r (right).

**Supplementary Figure 5.**
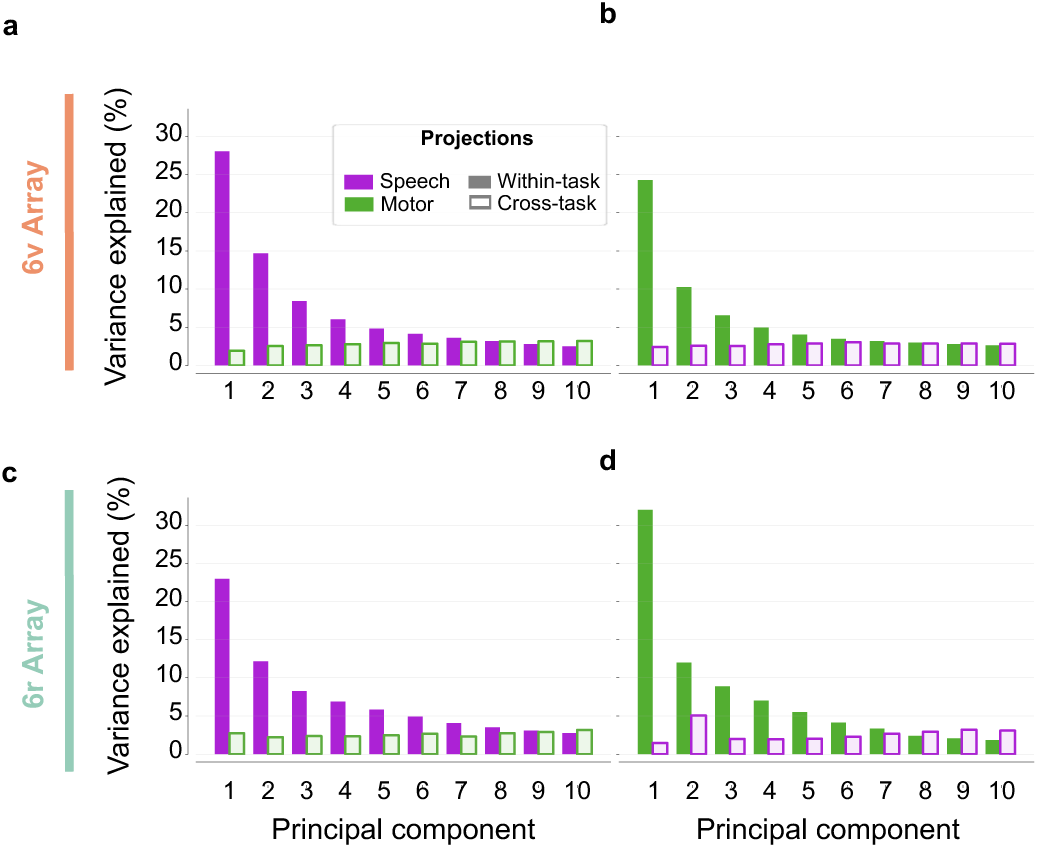
Variance captured by within- versus cross-task projections by individual principal components. This figure breaks down the cumulative variance capture shown in Fig. 6c by principal component. Bars reflect the variance captured when projecting neural activity onto each task-derived subspace axis (i.e., principal component) obtained via principal component analysis. “Within-task” refers to projecting each task’s activity onto its own subspace axes; “cross-task” refers to projecting one task’s activity onto the other task’s subspace axes. **(a)** Variance capture by projecting speech activity (purple bars), versus grasp activity (hollow green bars) onto speech subspace axes in 6v. **(a)** Variance capture by projecting grasp activity (green bars), versus speech activity (hollow purple bars) onto grasp subspace axes in 6v. **(c, d)** Same as **(a, b)**, respectively, for 6r.

**Supplementary Figure 6.**
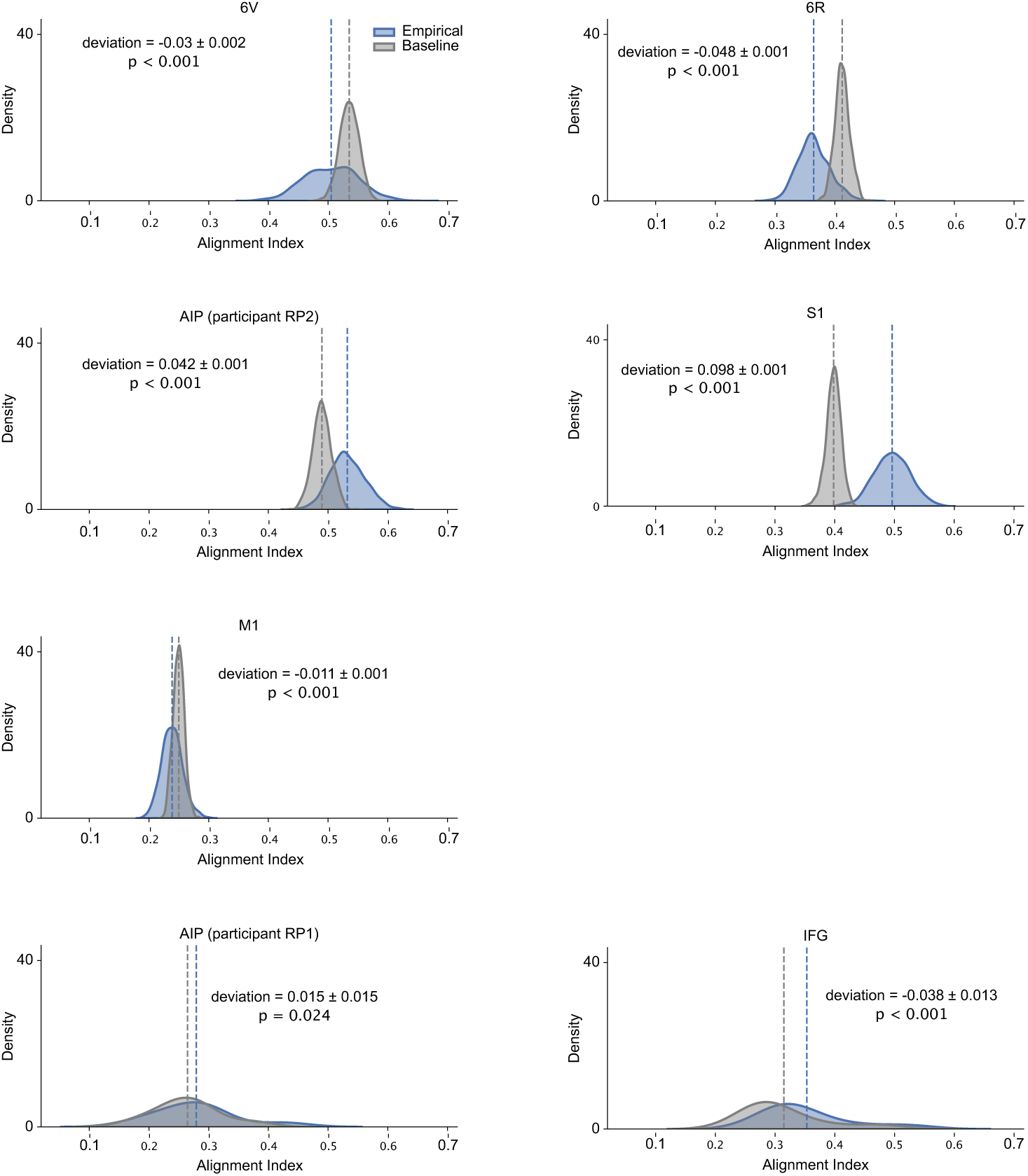
Cross-task subspace alignment distributions. Density plots showing the distribution of subspace alignment indexes for each array, providing an alternative view of the results in Fig. 6d. Alignment indexes range from 0 to 1, with higher values indicating greater overlap between speech and grasp subspaces. Significance was assessed by quantifying the deviation of empirical alignment indexes from baseline, where the baseline was obtained by projecting neural data onto randomly sampled subspaces within the same neural state space. Deviation is defined as empirical − baseline. Results are shown separately for each array.

